# Reconstruction of Fur pan-regulon uncovers the complexity and diversity of transcriptional regulation in *E. coli*

**DOI:** 10.1101/2020.05.21.109694

**Authors:** Ye Gao, Ina Bang, Yara Seif, Gayoung Nam, Anand V. Sastry, Ke Chen, Jonathan M. Monk, Kumari Sonal Choudhary, Sang Woo Seo, Eun-Yeol Lee, Donghyuk Kim, Bernhard O. Palsson

## Abstract

Regulons for many transcription factors have been elucidated in model strains leading to an understanding of their role in producing physiological states. Comparative analysis of a regulon and its target genes between different strains of the same species is lacking. Ferric uptake regulator (Fur), involved in iron homeostasis, is one of the most conserved TFs, and is present in a wide range of bacteria. Using ChIP-exo experiments, we performed a comprehensive study of Fur binding sites in nine *Escherichia coli* strains with different lifestyles. 79 of the 431 target genes (18%) found belong to Fur core regulon, comprising genes involved in ion transport and metabolism, energy production and conversion, and amino acid metabolism and transport. 179 of the target genes (42%) comprise the accessory regulon, most of which were related to cell wall structure and biogenesis, and virulence factor pathways. The remaining target genes (173 or 40%) were in the unique regulon, with gene functions that were largely unknown. Furthermore, deletion of the *fur* gene led to distinct phenotypes in growth, motility, antibiotic resistance, and the change of siderophore production. These results provide a more complete understanding of how Fur regulates a set of target genes with surprising variation in closely related bacteria.

## Introduction

Transcription factors (TFs) control gene expression by direct or indirect activation or repression of target genes. The binding state of TFs depends on the environmental conditions, such as presence or absence of effectors, and other physical stimuli ^1^. Many global TFs are highly conserved, which play defining physiological roles in a wide range of bacteria, enabling them to adapt to changing environments^2^. For example, iron is a critical cofactor for many enzymes in the bacteria. To maintain iron homeostasis, the arms race between the bacteria and their surrounding environments leads to the development of a sophisticated regulation network to control iron uptake. Ferric uptake regulator (Fur) is a major transcription factor to maintain iron homeostasis in the bacteria. Evidence suggests that Fur is involved in many other biological processes in different *E. coli* strains, including oxidative stress response, anaerobic metabolism, and even the infection of host cells^3^.

Next generation sequencing technologies have generated more than 10,000 *E. coli* genomes ^4^, which expands the scale of genome analysis from a few to hundreds of genomes. Meanwhile, pan-genome analysis becomes a reliable approach to estimate genome diversity in different strains^5^. Although Fur plays distinct roles in different *E. coli* strains, its protein sequence is broadly conserved from commensals to pathogens. The mechanism with which conserved TFs differentially affect bacterial physiological processes in closely related strains has been proposed ^6^. To understand the details underlying this mechanism, we should investigate the regulon, consisting of Fur, and target genes with their associated biological processes.

The approaches to estimate the regulon system were mostly derived from: 1) computational analysis of meta-genomics; 2) evolution of transcription factors, and 3) gene expression ^7^. But these approaches may not precisely reflect the regulon system, including the size and function of target genes. To investigate the regulon across different strains, it is required to reconstruct pan-regulon with the experimental data about how Fur regulates target genes in diverse *E. coli* strains. Pan-regulon is derived from pan-genome analysis, which consists of the entire set of target genes for all strains within one species. Similarly, it provides a framework for describing the diversity of regulon across different strains.

In this study, we combined pan-genome analysis with genome-scale experiments to reconstruct the strain-specific regulons (a set of genes directly controlled by a single TF), and pan-regulons (a total set of regulons from all the strains). To broaden our understanding of the regulation of conserved Fur, we generated the landscape of genome-wide binding sites of Fur across nine different *E. coli*, which are from the phylogroup A, B1, B2, D, and E. The results indicated 1672 genome-wide binding sites of Fur at iron-replete and iron-depleted conditions from nine different *E. coli* strains. By analyzing and comparing the genomes, we identified Fur regulons for each strain at iron-replete condition, and categorized regulons into three different groups: 1) core regulon: target genes are common to all of strains; 2) accessory regulon: target genes are found in more than one strain, but not all of them; and 3) unique regulon: target genes are found in only one strain. They consist of Fur pan-regulons in different *E. coli* strains, which are responsible for the metabolic pathways for iron uptake, storage, and energy production and conversion processes. Furthermore, virulence factor-related regulons showed remarkable differences between commensal and pathogenic strains.

Next, we systematically evaluated the functional differentiation of Fur between wild type and *Δfur* strains, especially for iron homeostasis, motility, and antibiotic sensitivity. In all, the results from nine *E. coli* strains, spanning all phylogenetic groups, reveal a pan transcriptional regulatory network (TRN) for Fur, and provide insights into understanding the diversity of regulation networks in multiple strains through pan-regulon.

## Materials & Methods

### Bacterial strains, media, and growth conditions

All strains (wild type, myc-tagged, and knockout strains) used in this study are listed in Dataset S1. For ChIP-exo experiments, the strains harboring 8-myc were generated by a λ red-mediated site-specific recombination system targeting C-terminal region as described previously^8^. For expression profiling by RNA-seq, *Δfur* strains from all strains were also constructed by a λ red-mediated site-specific recombination system^9^. For ChIP-exo experiments, glycerol stocks of 8-myc strains were inoculated into M9 minimal media (47.8 mM Na_2_HPO_4_, 22 mM KH_2_PO_4_, 8.6 mM NaCl, 18.7 mM NH_4_Cl, 2 mM MgSO_4_, and 0.1 mM CaCl_2_) with 0.2% (w/v) glucose. For iron-replete condition, M9 minimal media was supplemented with 0.1 mM FeCl_2_; for iron-depleted condition, M9 minimal media was supplemented with 0.2 mM 2,2’-dipyridyl at early-log phase and continue to culture at 37 °C for additional 2h with vigorous agitation; for regulatory condition, M9 minimal media was supplemented with 1 mL trace element solution (100X) containing 1 g EDTA, 29 mg ZnSO_4_.7H_2_O, 198 mg MnCl_2_.4H_2_O, 254 mg CoCl_2_.6H_2_O, 13.4 mg CuCl_2_, and 147 mg CaCl_2_. The culture was incubated at 37 °C overnight with agitation and then was used to inoculate the fresh media (1/200 dilution). The volume of the fresh media was 150 mL for each biological replicate. The fresh culture was incubated at 37 °C with agitation to the mid-log phase (OD_600_ ≈ 0.5). For RNA-seq expression profiling, glycerol stocks of all strains were inoculated into M9 minimal media with the same carbon sources as used in the ChIP-exo experiment. The concentration of carbon sources was 0.2% (w/v). For iron-replete condition, M9 minimal media was also supplemented with 0.1 mM FeCl_2_. The culture was incubated at 37 °C overnight with agitation and then was used to inoculate the fresh media. The fresh culture was incubated at 37 °C with agitation to the mid-log phase (OD_600_ ≈ 0.5).

### Bacterial motility assay

*E. coli* wild-type strains and *Δfur* strains were grown overnight at 37 ℃ on LB plates. Inoculation of a single colony onto motility plates (tryptone 1%, NaCl 0.25%, agar 0.3% and when indicated glucose was added) was done by using a toothpick. The motility halos were measured at 24 h. Each strain was evaluated in triplicate from independent plates.

### Measurement of bacterial growth

The effects of iron-limited conditions on cell growth were examined by growing all strains and *Δfur* strain under M9 minimal glucose medium without trace element solution. Cells grown overnight on M9 minimal glucose medium at 37 °C with agitation were inoculated into the fresh media, then were incubated at 37 °C with agitation. Similarly, to measure growth on iron-replete condition, the culture was incubated at M9 minimal media was supplemented with 0.1 mM FeCl_2_ at 37 °C overnight with agitation and then was used to inoculate the fresh media (1/200 dilution). The volume of the fresh media was 150 mL. The fresh culture was incubated at 37 °C with agitation. All of the growth curves were measured by three independent experiments at least and recorded by OD_600_ using Thermo BIOMATE 3S UV-visible spectrophotometer. The growth rate was calculated with GrowthRates 2.0^10^. The significant difference between wild type and *Δfur* strain was determined by the Student’s t test, P<0.01.

### ChIP-exo experiments

ChIP-exo experimentation was performed following the procedures as below. In brief, to identify Fur binding maps for each strain *in vivo*, the DNA bound to Fur from formaldehyde cross-linked cells were isolated by chromatin immunoprecipitation (ChIP) with the specific antibodies that specifically recognize myc tag (9E10, Santa Cruz Biotechnology), and Dynabeads Pan Mouse IgG magnetic beads (Invitrogen) followed by stringent washings as described previously^11^. ChIP materials (chromatin-beads) were used to perform on-bead enzymatic reactions of the ChIP-exo method^12^. Briefly, the sheared DNA of chromatin-beads was repaired by the NEBNext End Repair Module (New England Biolabs) followed by the addition of a single dA overhang and ligation of the first adaptor (5’-phosphorylated) using dA-Tailing Module (New England Biolabs) and NEBNext Quick Ligation Module (New England Biolabs), respectively. Nick repair was performed by using PreCR Repair Mix (New England Biolabs). Lambda exonuclease- and RecJf exonuclease-treated chromatin was eluted from the beads and overnight incubation at 65 °C reversed the protein-DNA cross-link. RNAs- and Proteins-removed DNA samples were used to perform primer extension and second adaptor ligation with following modifications. The DNA samples incubated for primer extension as described previously^13^ were treated with dA-Tailing Module (New England Biolabs) and NEBNext Quick Ligation Module (New England Biolabs) for second adaptor ligation. The DNA sample purified by GeneRead Size Selection Kit (Qiagen) was enriched by polymerase chain reaction (PCR) using Phusion High-Fidelity DNA Polymerase (New England Biolabs). The amplified DNA samples were purified again by GeneRead Size Selection Kit (Qiagen) and quantified using Qubit dsDNA HS Assay Kit (Life Technologies). Quality of the DNA sample was checked by running Agilent High Sensitivity DNA Kit using Agilent 2100 Bioanalyzer (Agilent) before sequenced using HiSeq 2500 (Illumina) following the manufacturer’s instructions. Each modified step was also performed following the manufacturer’s instructions. ChIP-exo experiments were performed in biological duplicates.

### RNA-seq expression profiling

Three milliliters of cells from mid-log phase cultures were mixed with 6 mL RNAprotect Bacteria Reagent (Qiagen). Samples were mixed immediately by vortexing for 5 s, incubated for 5 min at room temperature, and then centrifuged at 5000*g* for 10 min. The supernatant was decanted and any residual supernatant was removed by inverting the tube once onto a paper towel. Total RNA samples were then isolated using RNeasy Plus Mini kit (Qiagen) following the manufacturer’s instructions. Samples were then quantified using a NanoDrop 1000 spectrophotometer (Thermo Scientific) and quality of the isolated RNA was checked by running RNA 6000 Pico Kit using Agilent 2100 Bioanalyzer (Agilent). Paired-end, strand-specific RNA-seq library was prepared using KAPA RNA Hyper Prep kit (KAPA Biosystems), following the instruction ^14,1515^. Resulting libraries were analyzed on an Agilent Bioanalyzer DNA 1000 chip (Agilent). Sequencing was performed on a Hiseq 2500.

### Phylogenetic tree construction

A total of 401 complete genomic sequences of *E. coli* were downloaded from the Pathosystems Resource Integration Center database ^16^ and re-annotated using Prokka ^17^. The *E. coli* pan genome was built as in ^18^ using CD-HIT ^19^. Briefly, annotated coding DNA sequences were clustered into orthologous groups based on sequence similarity and the pan genome matrix was built in which rows represent orthologous groups, columns represent strains, and cell entries indicated whether an orthologous group was shared in each strain. In order to reduce the set of strains while maintaining the maximum phylogenetic diversity, strains were clustered into 40 groups (10% of samples) by running the scipy k-means algorithm minimizing the Euclidean distance between genome content. The pan genome matrix was used as input, and one representative strain was randomly chosen for each cluster. Each representative was then assigned to a multi-locus sequence type using the mlst GitHub repository which pulls its content from PubMLST ^20,21^, and to a phylogroup using ClermonTyping ^22^. Finally, we identified the orthologous groups which are conserved across all strains (core genes) but not duplicated in any of the selected strains. We concatenated the coding DNA sequences of the core genes for each strain and aligned the core genome using Parsnp ^23^. Recombination events were filtered out using Gubbins ^24^, and the phylogenomics tree was visualized in R using GGTREE ^25^.

### Peak calling for ChIP-exo dataset

Each read in raw sequencing data was trimmed to 31bp by using FASTX-toolkit^26^. The sequencing reads were mapped on each reference genome (Dataset S1) using Bowtie^27^. The SAM output files generated by Bowtie were changed to BAM format using SAMtools^28^. The peaks were predicted by the MACE program and were directly curated by Metascope visualization^29^. Each peak was adjusted to the 12-13bp size,which is already known in other paper. The calculation of S/N ratio resembles the way to calculate ChIP-chip peak intensity where IP signal was divided by Mock signal (https://sites.google.com/view/systemskimlab/software?authuser=0)

### Motif search from ChIP-exo peaks

The sequence motif analysis for Fur binding sites was performed using the MEME software suite^30^. For each strain, sequences in binding regions were extracted from the reference genome (Dataset S1). To achieve a more accurate motif, the sequence of each binding site was extended by 10bp at each end. The width parameter was fixed at 20bp and the minsites parameter was fixed at 90% of the total number of the sequence. All other parameters followed the default setting.

### Calculation of differentially expressed gene

The raw sequence reads of the RNAseq results were mapped onto each reference genome (Dataset S1) using Bowtie with the maximum insert size of 2000bp, and two maximum mismatches after trimming 3 bp at the 3’ ends^27^. SAM output files generated by Bowtie were changed to BAM format using SAMtools. We analyzed the differential gene expression using the DESeq package^31^. After the mapping, we count the number of reads that overlap each gene in the gff file, which contains the annotation information of each gene. Fragments per kilobase of exon per million fragments(FPKM) value for each gene was calculated. By comparing the WT and Fur-K/O condition, we could get the degree of expression change level. The differentially expressed genes were defined as genes with expression value with log_2_ (fold change) ≥ 1and *p*-value ≤ 0.05 or log2(fold change) ≤ −1.5 and *p*-value ≤ 0.05.

### Pan-regulon assembly and functional characterization

Using the Metascope program, we visualized each peak and annotation information in each strain. Some candidate operons were found for each peak, and they were specified as Fur-regulon according to their promoter location, presence of differential expression, and distance from the peak (Dataset S4). To confirm the co-transcriptional gene, we used the information from our RNAseq mapping result in WT strain, along with the information from EcoCyc, BioCyc, and DOOR data base^32,33,34^. For the pan-level analysis, we used the input file that includes all amino acid sequences of the specified Fur-regulon. BPGA options were identical to the options used for Pan-genome analysis. As above, the functional RNAs and pseudogenes were excluded from this analysis. Genes were grouped into orthologous gene families based on sequence similarity identified through the protein based Basic Local Alignment Search Tool using a cutoff of 80% percentage identity and a maximum e-val of 10^−6^ ^35^. Gene families were then functionally annotated and assigned to clusters of orthologous groups ontology using eggNOG ^36^. Genes were categorized into core, accessory and unique Fur regulon when they were regulated across all strains (core), a subset of strains (accessory), or one strain only (unique). They were further categorized as core, accessory and unique genomes when they were present in all strains (core), a subset of strains (accessory), or one strain only (unique).

### Virulence factor (VF) analysis

Virulence factors were annotated by searching for amino acid sequence similarity against the curated set of genes extracted from the virulence factor database (VFDB) ^37^. Virulence factors were categorized based on the description of each curated gene into heme uptake, siderophore, transport, fimbriae, AAI/SCI-II and other. By performing one-tailed Fisher’s exact test (Hypergeometric test) for virulence factor in Core, Accessory, and Unique category, the degree of enrichment was compared, and P-value <0.05 was considered significant.

### Clusters of Orthologous groups (COGs) enrichment

Fur regulons were categorized according to their annotated COG database^38^. Functional groups in core, accessory, and unique Fur-regulated genes were determined by COG categories.

## Results

### Characterization of ferric uptake regulator in diverse *E. coli* strains

Transcription factors (TFs) exhibit differential conservation in bacteria, when microorganisms respond and adapt to varying environments ^39^. However, many global TFs are still highly conserved in closely related bacteria. For example, the ferric uptake regulator (Fur) was broadly conserved orthologous transcription factors in *E. coli,* which is a master regulator to maintain iron homeostasis in the *E. coli ^13^*. To further investigate how diverse *E. coli* strains regulate their Fur transcriptional regulons, we developed the pan-regulon pipeline, to compare transcription factors and target genes together with their associated biological processes in diverse *E. coli* strains (Supplementary figure 1). The ferric uptake Pan-genome analysis allows us to analyze hundreds or thousands of genomes simultaneously. However, the scale of experimental verification is still limited to a few *E. coli* strains. Thus we should choose representative *E. coli* strains from different phylogroups.

To assess the diversity of *E. coli* strains, a total of 39 genomes were selected and reconstructed in a phylogenetic tree using a comparative genomics approach (Figure 1A). All of the strains consist of a variety of commensal and pathogenic strains, assigned to different phylogenetic groups: A, B1, B2, C, D, E, F, and G. Virulent strains mainly belong to B2, D, and E groups. In this study, a set of nine different strains were selected from the major phylogenetic groups: A, B1, B2, C, D, and E. They include: non-pathogenic strains (*E. coli* K-12 MG1655, W3110, BL21, Crooks, W, and KO11FL), and pathogenic strains (*E. coli* O44:H18 042, O6:H1 CFT073, and O157:H7 sakai belong to enteroaggregative *E. coli* (EAEC), uropathogenic *E. coli* (UPEC), and enterohemorrhagic *E. coli* (EHEC), respectively). The phylogenetic tree not only showed the evolutionary distance, but also indicated that these nine strains represent the diversity of E. *coli* strains in the phylogroup.

**Figure 1.**
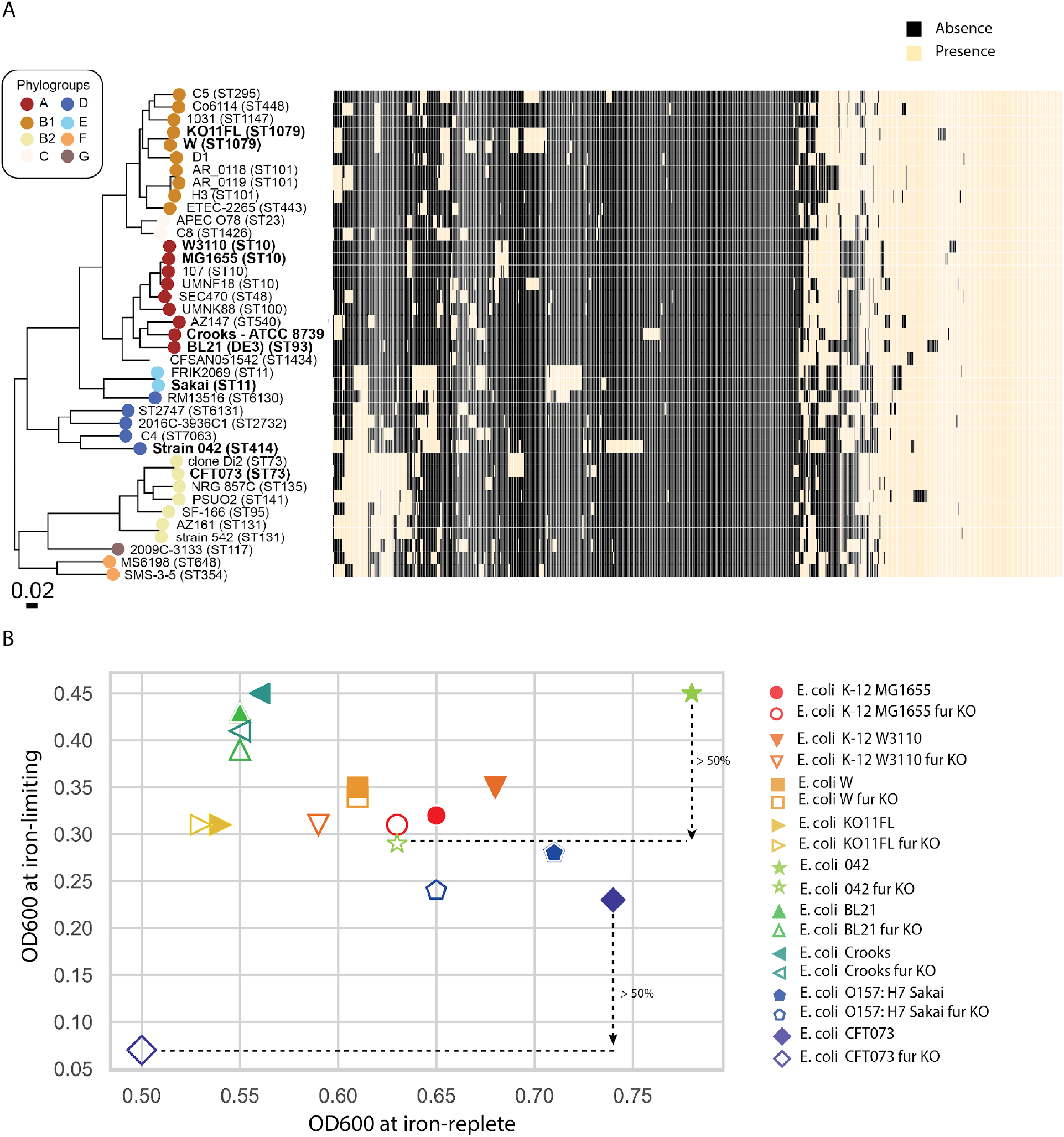
Characterization of ferric uptake regulator in diverse *E. coli* strains. **(A)** The phylogenetic tree of diverse *Escherichia coli* strains. Left panel: the tree lists 39 different *E. coli* strains. Both multilocus sequence types and phylogroups are portrayed. Each phylogroup (A, B1, B2, C, D, E, F, G) was marked by specific color. The nine representative strains used in this study are in bold. The phylogeny was constructed by running the scipy k-means algorithm minimizing the Euclidean distance between genome content, and visualized in R using GGTREE. Right panel: visualization of pan-regulon among 39 different *E. coli* strains using the matrix. (**B**) The final OD_600_ of nine *E. coli* representative strains at iron-replete and iron-limiting condition, respectively. X-axis: OD600 at iron-replete condition. The cells grow at M9 minimal medium with 0.1 mM FeCl_2_. Y-axis: OD600 at iron-limiting condition. The cells grow at M9 minimal medium without any iron source. Absorbance at 600 nm was measured every 15 min using a TECAN Infinite 200 automated microbiology growth analysis system.

To confirm the conservation of Fur, the protein sequence alignment was examined across these representative strains (Supplementary figure 2), which showed that all of the amino acids in Fur protein are 100% identical across these strains, though their genome sequences vary. These information raised an important question, how broadly conversed Fur regulates target genes to play distinct functions in different *E. coli* strains.

Furthermore, to investigate the effects of Fur on the physiological roles in *E. coli*, aerobic growth in iron-replete and iron-limiting conditions was analyzed. All of the strains were capable of growing under both conditions (Figure 1B). There were no significant differences with or without ferrous in *E. coli* MG1655, W3110, Crooks, W, KO11FL, BL21, or Sakai (t-test, P > 0.05). However, *E. coli* CFT073 and 042 showed significant differences at final OD_600_, which might be attributed to the capacity of iron uptake (t-test, P < 0.05). Overall, this result demonstrated that deletion of Fur, involved in iron homeostasis, led to defective growth in most pathogenic strains.

Next, to explore strain-specific diversity of the fur regulon, the myc-tagging strains were constructed for these representative strains except for *E. coli* MG1655.

### The comparison of genome-wide Fur binding sites across the *E. coli* strains

To reconstruct the pan regulon, we combined Fur ChIPexo with gene expression profiling data to identify the strain-specific Fur regulon, and then generate pan regulon across *E. coli* strains (Figure 2A). Fur-binding sites in *E. coli* K-12 MG1655 have been characterized by *in vivo* experiments ^13^. Here,we focused on the ChIP-exo and RNAseq results at iron-replete conditions. Similarly, we utilized ChIP-exo to explore the binding sites of Fur at iron-deplete condition as well (Supplementary figure 3).

**Figure 2.**
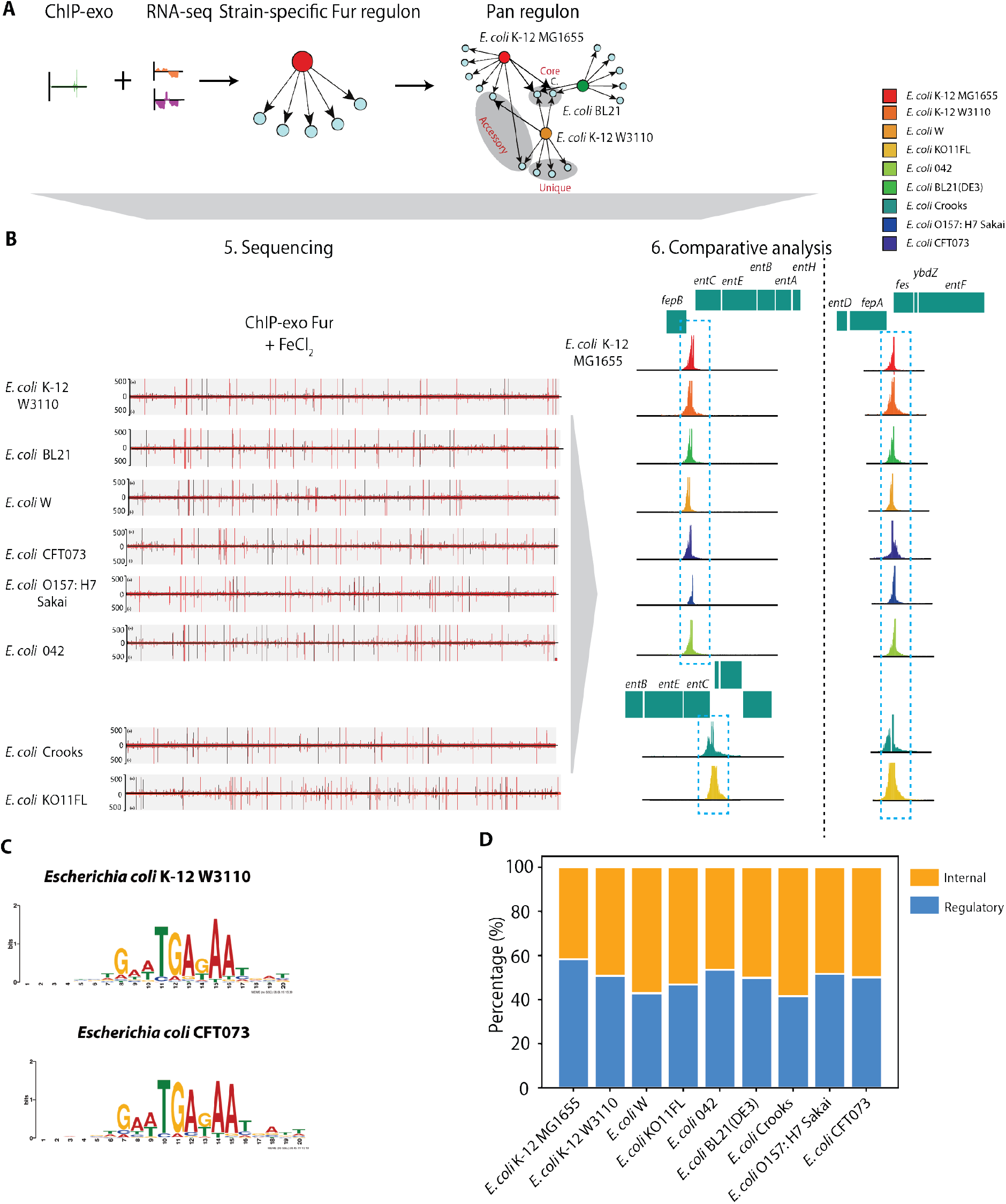
Comparison of genome-wide binding of Fur across different *E. coli* strains. (A) The pipeline to reconstruct pan-regulon across different E. coli strains **(B)** Comparison of genome-wide binding in nine *E. coil* strains at iron-replete condition; **(C)** The representative Fur boxes (consensus sequences) from commensal (*E. coli* W3110) and pathogenic *E. coli* (*E. coli* CFT073). **(D)** The percentage of binding peaks distributed at regulatory and internal regions across different strains.

Using the peak calling algorithm MACE, a total of 1322 reproducible Fur-binding peaks were identified across these representative strains (Figure 2B). Interestingly, the genome-wide bindings of Fur showed that they have similar binding numbers, binding motifs, and distribution of binding peaks, thus Fur binding sites are conserved across *E. coli* strains. First, the number of binding events from *E. coli* were very similar except for *E. coli* ATCC 55124 (KO11FL). Second, most of the binding peaks observed at iron starvation were overlapped with iron-replete conditions, though the overlapped percentage varies among the strains (Supplementary figure 5). As previous studies demonstrated, Fur has two conformations, apo- and holo-Fur, which depends on the availability of iron. In iron-replete condition, Fur is activated by Fe^2+^, and formed as holo-Fur, which represses target genes involved in the iron acquisition system. Thus, there are four different modes of Fur regulation *in vivo*, including holo-Fur repression (HR), holo-Fur activation (HA), apo-Fur repression (AR), and apo-Fur activation (AA). It is possible that apo-Fur has distinct target genes, though the function of Fur only occupied at iron starvation condition is still unclear. Third, the enrichment fold of Fur binding at iron-replete condition is higher than that at iron starvation condition (Supplementary figure 6). Also, Fur-box (DNA-binding motif) is broadly conserved across different strains. Two representative motifs from W3110 (non-pathogenic strain) and CFT073 (pathogenic strain) were listed (Figure 2C). The rest of them were listed in supplementary figure 7. Another similarity was about the percentage of Fur binding sites within putative regulatory regions (promoters and 5’-proximal to coding regions) (Figure 3D, and supplementary figure 8). Overall, the observation from genome-wide binding showed that conserved Fur has broadly similar binding features across *E. coli* strains.

**Figure 3.**
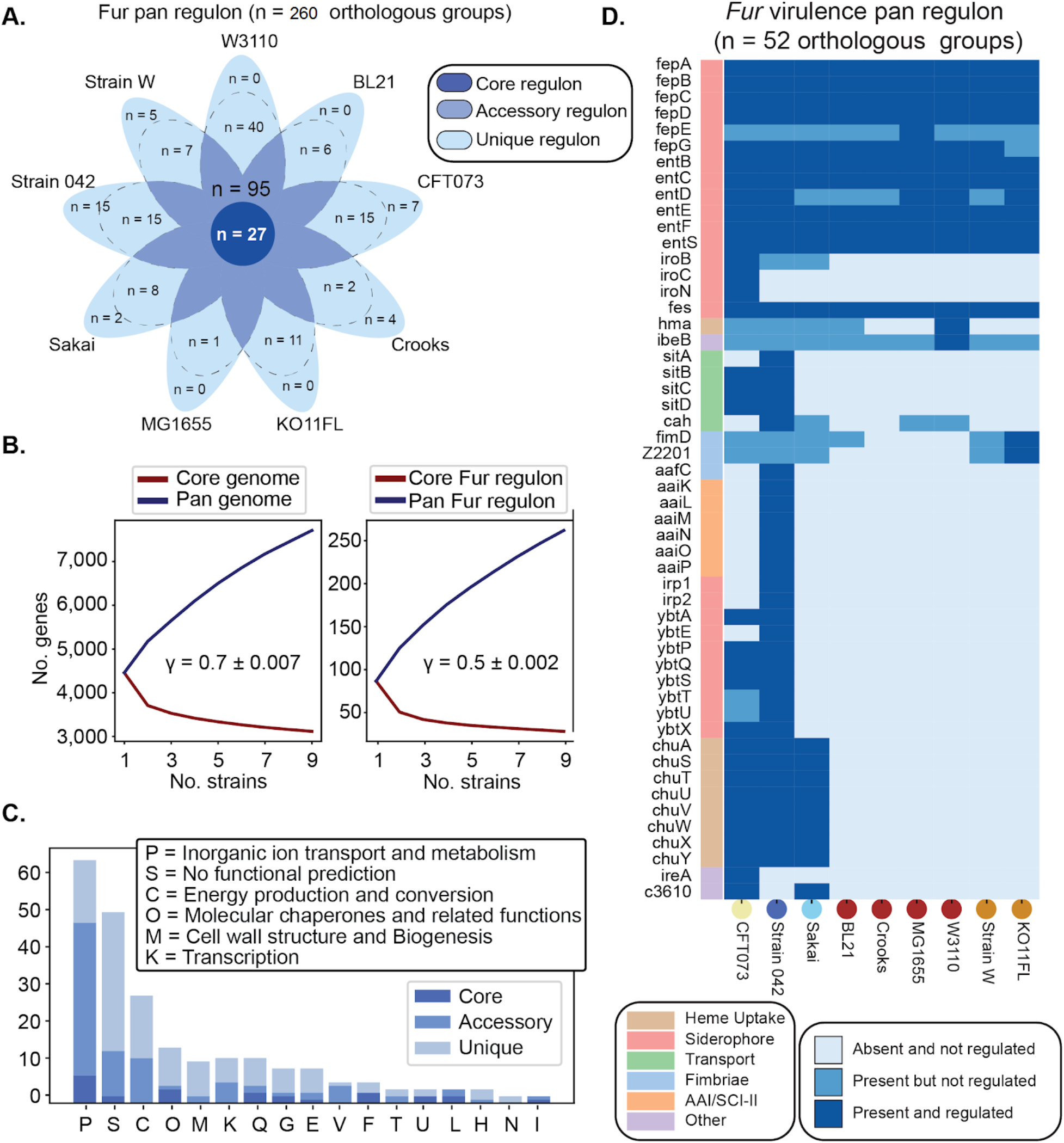
The pan-regulon of Fur across nine *E. coli* strains. (A) Comparison of the strain-specific Fur regulons. The venn diagram shows the distribution of orthologous groups that are regulated by Fur across nine *E. coli* strains into the core, accessory and unique fur regulon. The unique regulon contains genes that are present across multiple strains. As such, we display two numbers for the unique regulon separated by a dashed line: 1) the number of genes that are uniquely carried in one strain, and uniquely regulated in one strain (displayed outside of the dashed lines), and; 2) the number of genes that are part of the unique fur regulon but are shared across multiple strains. (B) Pan and core genome curve, and pan and core Fur regulon curves for the nine *E. coli* strains. Strains were randomly sampled and drawn one at a time. We drew the pan genome and pan regulon curves as in ^18^. Briefly, strains were randomly sampled, and drawn one at a time. The number of novel genes found at each draw is added to the pan curve, and the number of genes that are conserved across all draws is retained for the core curve. The gene sets considered for the pan genome are the totality of the genes found in each genome, and the gene sets considered for the pan regulon are the genes which were experimentally determined to be regulated by Fur (see section Pan-regulon assembly and functional characterization). Heaps’ law n = k N-(1 -γ)was fit to the pan genome and pan regulon curves to yield decay rates (= 1 -γ) for the pan genome and pan regulon curves. When γ> 0, the pan genome is considered to be “open”, meaning that the number of new genes discovered as more strains are analysed is not capped. (C) Distribution of Fur pan-regulon genes per functional category as determined by the clusters of orthologous groups ontology (COGs). The inorganic ion transport and metabolism and the energy production and conversion were the least conserved categories. (D) Distribution of virulence factors of the Fur pan regulon. The two most represented categories are siderophore production and heme uptake. Strains CFT073 and 042 have the largest numbers of virulence factors regulated by Fur. Some virulence factors such as *hma* are regulated by Fur in one strain (W3110) but not in other strains (CFT073, strain 042, Sakai and BL21), despite being present in these strains.

Surprisingly, we observed one outlier strain, *E. coli* KO11FL, which has the largest number of Fur-binding events. To demonstrate the relationship between genome size and binding peaks, the correlation between them was analyzed (Supplementary figure 4). *E. coli* KO11FL strain is a current lab version of KO11 that was engineered from parental strains of *E. coli W* (ATCC 9637). Compared to the parental W strain, KO11FL has extensive chromosomal rearrangement, and tandem repeat region, consisting of at least 20 copies of 10-kb unit ^40^. It is likely that the genome rearrangement caused more Fur binding events.

### Reconstruction of strain-specific Fur regulon and pan regulon in *E. coli* strains

To reconstruct the fur regulons, the gene *fur* was knocked out by using lambda recombination in all strains. The expression profiling data from each wild type and *Δfur* strain was analyzed at iron-replete condition, thus we focused on strain-specific Fur regulon and pan regulon at iron-replete condition in this study.

Combining ChIP-exo binding sites with expression data, we found that the Fur pan regulon contains a total of 260 orthologous groups, of which 65 are involved in inorganic ion transport and metabolism, and 64 have an unknown function. Next, we subdivided the Fur pan regulon into: 1) the core regulon (n = 27 target genes regulated by Fur across all strains), 2) the accessory regulon (n = 95 target genes regulated in a subset of strains), and 3) the unique regulon (n = 138 target genes uniquely regulated in only one strain) (Figure 3A). On average, only 40% of the fur regulon in a strain is part of the core fur regulon. Surprisingly, regulation does not reflect the level of sequence conservation across strains. In total, 117 (40%) orthologous groups were conserved, but only 27 were regulated across all strains. These results reveal that the fur regulatory network changes on a strain-to-strain basis, regardless of the functional gene content.

To evaluate the relationship between the number of strains and the size of pan-genome, pan and core genome curves for the nine *E. coli* strains were constructed (Figure 3B, left panel). Similarly, pan and core Fur regulon curves were shown at the right panel (Figure 3B). Heaps’ law n = k N-(1 -γ) was fit to the pan genome and pan regulon curves to yield their decay rates (= 1 -γ) γ> 0, the pan genome is considered to be “open”, meaning that the number of new genes discovered as more strains are analysed is not capped.

Furthermore, the Fur pan regulon was distributed across 17 functional groups, with the most represented groups being inorganic ion transport and metabolism, unknown functions, and energy production and conversion (Figure 3C). The core regulon is enriched with inorganic ion transport and metabolism while the unique regulon is enriched with proteins with no functional prediction. In total, virulence factors constituted 20% (52 out of 260) of the fur pan regulon, of which only nine were commonly regulated by fur across all strains including fepABCD and entBCEFS (Figure 3D). Strains 042 (n = 40), CFT073 (n = 31), and Sakai (n = 19), have the largest numbers of fur regulated virulence factors among nine *E. coli* strains.

### Loss of Fur leads to distinct phenotypes in *E. coli*

Fur is involved in many other biological processes, including production of siderophore, motility, and antibiotic resistance. Here, we examined the phenotypes between wild type and *Δfur* strains as below.

First, we found that *Δfur* strains generated a high level of siderophore compared to the wide type, when cells grew to the stationary phase (Supplementary figure 9). Second, the influence of Fur has been studied in the flagella biosynthesis and motility in the pathogenic bacteria (non-*E. coli*) ^41, 42^. However, there is little information about the influence of Fur in different *E. coli* strains. Here we firstly examined if loss of Fur affects the flagella biosynthesis. The motility test showed that loss of Fur has no effect on *E. coli* flagella (Supplementary figure 10 and 11). Furthermore, we investigate the function of flagella on the low agar plates. As shown in supplementary figure 12, after 24 h incubation, three different types of motility patterns were presented among the different strains. i) *E. coli* MG1655, W3110, Crooks, and BL21, which belong to phylogroup A, all showed limited motility; ii) *E. coli* W and KO11FL, which belong to phylogroup B1, presented high capacity of motility with or without glucose; iii) *E. coli* CFT073 and Sakai, which belong to phylogroup B2 and E, showed reduced motility in the absence of glucose. But for *E. coli* 042, its motile behavior didn’t change too much with or without glucose.

These results pointed out that non-pathogenic and pathogenic strains showed different motility phenotypes in the planktonic stage, though loss of Fur didn’t affect the flagella biosynthesis. It is important to observe that the addition of glucose to the media may improve the motility for pathogenic strains. That might contribute to the adaptation of pathogenic strains to different environments or nutrient conditions. For example, the regulation of glucose catabolism and glycolysis was reported to be coupled to virulence factor expression in the EHEC strain (Sakai)^43^.

Next, we investigated the effect of Fur on the antibiotic sensitivity in *E. coli*. A previous study demonstrated that perturbation of iron homeostasis facilitated the evolution of ciprofloxacin resistance in *Escherichia coli* ^44^. In this study, we utilized the phenotype microarrays, and compared the responses between wild type and *Δfur* mutants. We observed that the loss of Fur regulon significantly increased sensitivity to at least one of the antibiotics (Figure 4A). Another bacterial adaptive study investigated the transcriptome profiles of *E. coli* upon adaptation to some antibiotics and biofuels, and found expression change of a set of genes (*tar, fliA, fiu, mntH, amtB, citC, entC, entE, wzc, yfiL, yjjZ*) associated with adaptive resistance^45^. We investigated Fur pan-regulons, and found that most of these genes reported belong to Fur regulons (Supplementary table 2). For example, fiu, mntH, entC, entE, yjjZ belong to core regulon, which were upregulated after deletion of Fur. Accordingly, we proposed the model for the mechanism of Fur regulation on the antibiotic response (Figure 4B).

**Figure 4.**
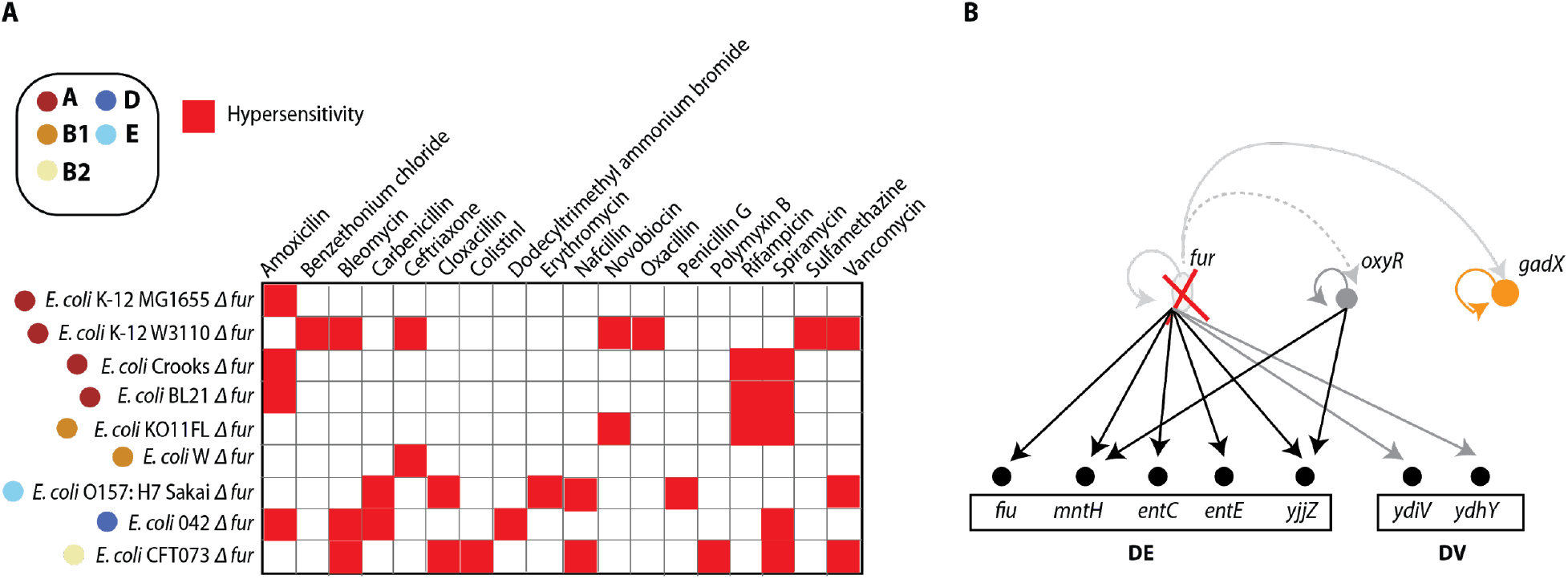
The antibiotic sensitivity profiles with *fur* deleted mutants from phenotype microarray tests, compared to each wild type strain. (A) The antibiotic responses profiles in *E. coli* strains; (B) The model for the mechanism of Fur regulation on the antibiotic response.

### Reconstruction of Fur extended “regulon” from genome-wide binding sites

In this study, we used the term “core regulon” to refer to target genes that are present in all bacteria considered here; “accessory regulon” refers to target genes that are found in more than one strain; and “unique regulon” refers to target genes that are found in only one strain. Also, “target genes” means the genes directly regulated by Fur, which combines genome-wide binding with the gene expression data in *Δfur* strains (|log_2_ (fold change)| > 1.5, and q-value < 0.05). But these traditional criteria are limited to a small range of target genes directly regulated by Fur.

To fully understand the mechanism of Fur differential regulation, we expanded the range of regulons to the genes with Fur boxes (genes with Fur binding sites at either promoter or intragenic region) (Supplementary table 3). Additionally, we tracked these regulons from their origin at the pan genome. For example, we observed that the gene (representative name: cluster_476) belongs to a unique regulon, though it is shared with all strains (core genome) (supplementary material Dataset 4). Because the fold change of expression level is below the cut-off value (|log 2| < 1.5) in all *Δfur* strains except for *E. coli* KO11FL. Similarly, another gene (cluster_1057) is classified into a unique regulon, though it is from the accessory genome. These results suggest that the expansion of regulons to all genes containing Fur-boxes may give us insights into how Fur differentially influences expression at the transcriptional level.

### The Fur transcriptional regulation varies across different *E. coli* strains

Nine *E. coli* strains were analyzed to comprehensively compare Fur strain-specific transcriptional regulation (Figure 5A). The relationships between each strain were used to examine whether or not Fur transcriptional regulation is conserved. First, *E. coli* K-12 W3110 has one of the highest conserved Fur regulation across these strains. Second, *E. coli* KO11FL has the lowest conservation compared to other strains. As discussed before, the chromosomal rearrangements and multiple tandem copies at the genome leads to disparate Fur binding events. Third, Fur transcriptional regulation in *E. coli* KO11FL differs most from that in pathogenic *E. coli* 042. These results demonstrated that Fur transcriptional regulation varies at the strain level.

**Figure 5.**
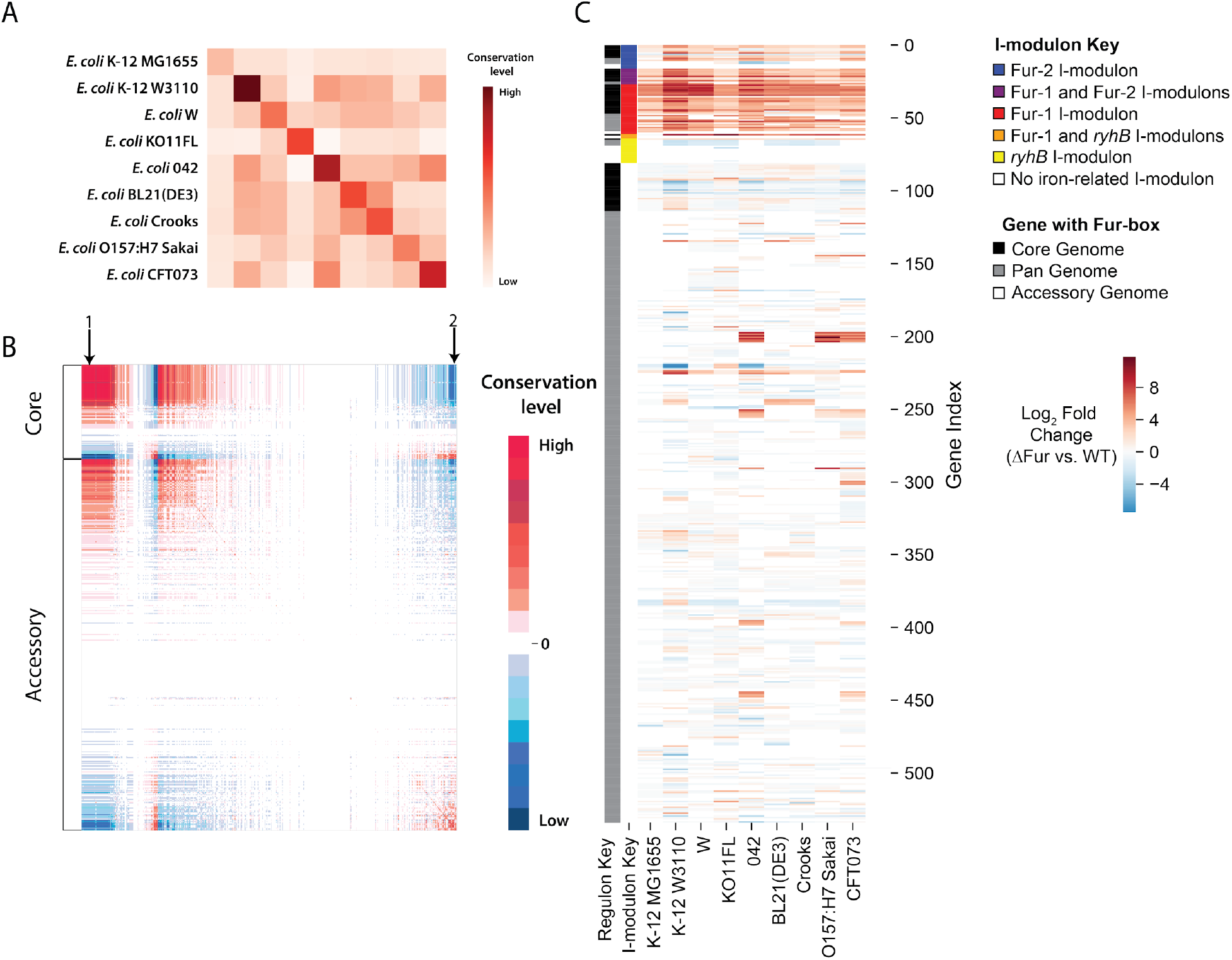
Conservation variation of strain-specific Fur transcriptional regulation. (A) Comparison of strain-specific Fur transcriptional regulation across different strains. Both raw and column index represent the strains in the same order. Each element in the matrix represents strain-specific Fur transcriptional regulation. The diagonal line is the relationship between themselves. (B) Comparison of genes containing Fur-box across different strains. The upper panel represents genes with Fur-box from the core genome; the bottom panel represents genes from the accessory genome.(C) The independent component analysis extracts regulatory modulons from pan-regulons and accessory regulons. Pan-regulons mainly consist of Fur-1, Fur-2, and RyhB i-modulons. The color bar corresponds to the value (log2 fold change between wild type and *Δfur* strains). The distribution of Fur-1, Fur-2, and RyhB i-modulons in the accessory regulons.

To further examine the influence of Fur at the transcriptional level, we quantitatively checked the regulation for all the genes with Fur-boxes. We observed that the highest conservation region contains core-regulon associated with iron homeostasis (arrow 1, Figure 5B). All of them were highly expressed after deletion of the repressor Fur. However, the genes related with virulence factors from the accessory genome show differentially regulation patterns (arrow 2, Figure 8B). These observations demonstrated that Fur transcriptional regulation varies across the pan-genome.

To compare Fur pan-regulon and i-modulons, these genes with Fur-boxes were examined using independent component analysis (ICA), which extraced Fur related i-modulons that overlay with pan-regulon^46^ (Figure 5C). That data showed that genes with Fur-box include most of Fur i-modulon (Fur-1, Fur-2, and ryhB i-modulon).

## Discussion

The details underlying the regulation of conserved TFs in closely related bacteria are unclear. To address this question, we combined comparative genomics (pan-genome) with high-throughput experimental approaches, to reconstruct pan-regulon (including core, accessory, and unique regulon) across phylogenetically related *E. coli* strains. Specifically, Fur was chosen as one of the highly conserved TFs in the *E. coli*. On one hand, its highly conserved sequence recognizes similar DNA target sequences in different strains; on the other hand, as a global transcriptional factor, it plays central roles in many aspects, including but not limited to iron homeostasis, redox oxidative stress response, motility, and pathogenesis, with some of which bacteria adapt to distinct environments. Meanwhile, nine *E. coli* strains, spanning distinct phylogroups (A, B1, B2, D, and E), represent a set of commensal and pathogenic ones in this species. In all, the details of how conserved Fur organizes its regulons at the genomes across different strains were explored in this study.

We developed a pipeline to reconstruct Fur regulon based on genome-wide binding sites and differentially expressed genes for each strain. Next, we categorized regulons into three groups: core, accessory, and unique regulons, based on pan-genome. The pan regulons presented multiple functions that Fur is involved in, and differentially expressed targets across different strains distinguish Fur regulation from each other. To further explore differential regulation, Fur regulons (refers to target genes regulated by Fur) were extended to all of the genes with Fur box at the genomes (so-called extended regulons). These results demonstrated how conserved TFs regulate a core set of genes, or strain-specific genes with which bacteria adapt to distinct habitats, and provided new insights into the biological functions of Fur regulons across different strains. These data suggested that the homologous regulons vary even among closely related species, possibly reflecting species-specific adaptation to environments.

## Data availability

The whole dataset of ChIP-exo and RNA-seq has been deposited to GEO with the accession number of GSE150240 and GSE150501.

## Supplementary data

Supplementary data are available online.

## Acknowledgement

We thank Richard Szubin for help with ChIP-exo and RNA-seq library sequencing. We thank Dr. Zachary A. King, Dr. Amitesh Anand, and Dr. Bin Du for helpful discussions. We thank Marc Abrams for reviewing and editing the manuscript.

## Author Contributions

Y.G., D.K. and B.O.P. designed the study. Y.G. performed experiments. Y.G., I.B., Y.S., G.N., A.V.S., J.M.M., K.S.C., S.W.S., E.Y.L., and D.K. analyzed data. Y.G., Y.S., K.C., D.K. and B.O.P. wrote the manuscript, with contributions from all other authors.

## Reference

1. Bervoets, I. & Charlier, D. Diversity, versatility and complexity of bacterial gene regulation mechanisms: opportunities and drawbacks for applications in synthetic biology. FEMS Microbiol. Rev. 43, 304–339 (2019).

2. Balleza, E. et al. Regulation by transcription factors in bacteria: beyond description. FEMS Microbiol. Rev. 33, 133–151 (2009).

3. Fillat, M. F. The FUR (ferric uptake regulator) superfamily: diversity and versatility of key transcriptional regulators. Arch. Biochem. Biophys. 546, 41–52 (2014).

4. Abram, K. et al. What can we learn from over 100,000 Escherichia coli genomes? doi:10.1101/708131.

5. Vernikos, G., Medini, D., Riley, D. R. & Tettelin, H. Ten years of pan-genome analyses. Curr. Opin. Microbiol. 23, 148–154 (2015).

6. Leclerc, J.-M., Dozois, C. M. & Daigle, F. Salmonella enterica serovar Typhi siderophore production is elevated and Fur inactivation causes cell filamentation and attenuation in macrophages. FEMS Microbiology Letters vol. 364 (2017).

7. Perez, J. C., Christian Perez, J. & Groisman, E. A. Evolution of Transcriptional Regulatory Circuits in Bacteria. Cell vol. 138 233–244 (2009).

8. Cho, B.-K., Knight, E. M. & Palsson, B. O. PCR-based tandem epitope tagging system for Escherichia coli genome engineering. Biotechniques 40, 67–72 (2006).

9. Datta, S., Costantino, N. & Court, D. L. A set of recombineering plasmids for gram-negative bacteria. Gene 379, 109–115 (2006).

10. Hall, B. G., Acar, H., Nandipati, A. & Barlow, M. Growth rates made easy. Mol. Biol. Evol. 31, 232–238 (2014).

11. Cho, B.-K., Kim, D., Knight, E. M., Zengler, K. & Palsson, B. O. Genome-scale reconstruction of the sigma factor network in Escherichia coli: topology and functional states. BMC Biol. 12, 4 (2014).

12. Rhee, H. S. & Pugh, B. F. ChIP-exo method for identifying genomic location of DNA-binding proteins with near-single-nucleotide accuracy. Curr. Protoc. Mol. Biol. Chapter 21, Unit 21.24 (2012).

13. Seo, S. W. et al. Deciphering Fur transcriptional regulatory network highlights its complex role beyond iron metabolism in Escherichia coli. Nat. Commun. 5, 4910 (2014).

14. Ross, M. G. et al. Characterizing and measuring bias in sequence data. Genome Biol. 14, R51 (2013).

15. Quail, M. A. et al. Optimal enzymes for amplifying sequencing libraries. Nat. Methods 9, 10 (2011).

16. Wattam, A. R. et al. Improvements to PATRIC, the all-bacterial Bioinformatics Database and Analysis Resource Center. Nucleic Acids Res. 45, D535–D542 (2017).

17. Seemann, T. Prokka: rapid prokaryotic genome annotation. Bioinformatics 30, 2068–2069 (2014).

18. Seif, Y. et al. Genome-scale metabolic reconstructions of multiple Salmonella strains reveal serovar-specific metabolic traits. Nat. Commun. 9, 3771 (2018).

19. Huang, Y., Niu, B., Gao, Y., Fu, L. & Li, W. CD-HIT Suite: a web server for clustering and comparing biological sequences. Bioinformatics 26, 680–682 (2010).

20. Jolley, K. A. & Maiden, M. C. J. BIGSdb: Scalable analysis of bacterial genome variation at the population level. BMC Bioinformatics 11, 595 (2010).

21. Seemann, T. mlst. (Github).

22. Beghain, J., Bridier-Nahmias, A., Le Nagard, H., Denamur, E. & Clermont, O. ClermonTyping: an easy-to-use and accurate in silico method for Escherichia genus strain phylotyping. Microb Genom 4, (2018).

23. Treangen, T. J., Ondov, B. D., Koren, S. & Phillippy, A. M. The Harvest suite for rapid core-genome alignment and visualization of thousands of intraspecific microbial genomes. Genome Biol. 15, 524 (2014).

24. Croucher, N. J. et al. Rapid phylogenetic analysis of large samples of recombinant bacterial whole genome sequences using Gubbins. Nucleic Acids Res. 43, e15 (2015).

25. Yu, G., Smith, D. K., Zhu, H., Guan, Y. & Lam, T. T.-Y. ggtree: an r package for visualization and annotation of phylogenetic trees with their covariates and other associated data. Methods Ecol. Evol. 8, 28–36 (2017).

26. Yee, L. L. W. & Tapani, T. M. Bioinformatics: A Practical Handbook Of Next Generation Sequencing And Its Applications. (#N/A, 2017).

27. Langmead, B., Trapnell, C., Pop, M. & Salzberg, S. L. Ultrafast and memory-efficient alignment of short DNA sequences to the human genome. Genome Biol. 10, R25 (2009).

28. Li, H. et al. The Sequence Alignment/Map format and SAMtools. Bioinformatics 25, 2078–2079 (2009).

29. Wang, L. et al. MACE: model based analysis of ChIP-exo. Nucleic Acids Res. 42, e156 (2014).

30. Bailey, T. L. et al. MEME SUITE: tools for motif discovery and searching. Nucleic Acids Res. 37, W202–8 (2009).

31. Anders, S., Anders, S. & Huber, W. Differential expression analysis for sequence count data. Nature Precedings (2010) doi:10.1038/npre.2010.4282.1.

32. Keseler, I. M. et al. The EcoCyc database: reflecting new knowledge about Escherichia coli K-12. Nucleic Acids Res. 45, D543–D550 (2017).

33. Karp, P. D. et al. The BioCyc collection of microbial genomes and metabolic pathways. Brief. Bioinform. (2017) doi:10.1093/bib/bbx085.

34. Mao, X. et al. DOOR 2.0: presenting operons and their functions through dynamic and integrated views. Nucleic Acids Res. 42, D654–9 (2014).

35. Altschul, S. F., Gish, W., Miller, W., Myers, E. W. & Lipman, D. J. Basic local alignment search tool. J. Mol. Biol. 215, 403–410 (1990).

36. Huerta-Cepas, J. et al. eggNOG 5.0: a hierarchical, functionally and phylogenetically annotated orthology resource based on 5090 organisms and 2502 viruses. Nucleic Acids Res. 47, D309–D314 (2019).

37. Liu, B., Zheng, D., Jin, Q., Chen, L. & Yang, J. VFDB 2019: a comparative pathogenomic platform with an interactive web interface. Nucleic Acids Res. 47, D687–D692 (2019).

38. Tatusov, R. L. et al. The COG database: an updated version includes eukaryotes. BMC Bioinformatics 4, 41 (2003).

39. Galán-Vásquez, E., Sánchez-Osorio, I. & Martínez-Antonio, A. Transcription Factors Exhibit Differential Conservation in Bacteria with Reduced Genomes. PLoS One 11, e0146901 (2016).

40. Turner, P. C. et al. Optical mapping and sequencing of the Escherichia coli KO11 genome reveal extensive chromosomal rearrangements, and multiple tandem copies of the Zymomonas mobilis pdc and adhB genes. J. Ind. Microbiol. Biotechnol. 39, 629–639 (2012).

41. Troxell, B. & Hassan, H. M. Transcriptional regulation by Ferric Uptake Regulator (Fur) in pathogenic bacteria. Front. Cell. Infect. Microbiol. 3, 59 (2013).

42. Tanui, C. K., Shyntum, D. Y., Priem, S. L., Theron, J. & Moleleki, L. N. Influence of the ferric uptake regulator (Fur) protein on pathogenicity in Pectobacterium carotovorum subsp. brasiliense. PLoS One 12, e0177647 (2017).

43. Njoroge, J. W., Nguyen, Y., Curtis, M. M., Moreira, C. G. & Sperandio, V. Virulence meets metabolism: Cra and KdpE gene regulation in enterohemorrhagic Escherichia coli. MBio 3, e00280–12 (2012).

44. Méhi, O. et al. Perturbation of iron homeostasis promotes the evolution of antibiotic resistance. Mol. Biol. Evol. 31, 2793–2804 (2014).

45. Erickson, K. E., Otoupal, P. B. & Chatterjee, A. Transcriptome-Level Signatures in Gene Expression and Gene Expression Variability during Bacterial Adaptive Evolution. mSphere 2, (2017).

46. Sastry, A. V. et al. The Escherichia coli Transcriptome Consists of Independently Regulated Modules. bioRxiv 620799 (2019) doi:10.1101/620799.

